# Rapid and assured genetic engineering methods applied to *Acinetobacter baylyi* ADP1 genome streamlining

**DOI:** 10.1101/754242

**Authors:** Gabriel A. Suárez, Kyle R. Dugan, Brian A. Renda, Sean P. Leonard, Lakshmi S. Gangavarapu, Jeffrey E. Barrick

## Abstract

One goal of synthetic biology is to improve the efficiency and predictability of living cells by removing extraneous genes from their genomes. We demonstrate improved methods for engineering the genome of the metabolically versatile and naturally transformable bacterium *Acinetobacter baylyi* ADP1 and apply them to a genome streamlining project. In Golden Transformation, linear DNA fragments constructed by Golden Gate Assembly are directly added to cells to create targeted deletions, edits, or additions to the chromosome. We tested the dispensability of 55 regions of the ADP1 chromosome using Golden Transformation. The 19 successful multiple-gene deletions ranged in size from 21 to 183 kilobases and collectively accounted for 24.6% of its genome. Deletion success could only be partially predicted on the basis of a single-gene knockout strain collection and a new Tn-Seq experiment. We further show that ADP1’s native CRISPR/Cas locus is active and can be retargeted using Golden Transformation. We reprogrammed it to create a CRISPR-Lock, which validates that a gene has been successfully removed from the chromosome and prevents it from being reacquired. These methods can be used together to implement combinatorial routes to further genome streamlining and for more rapid and assured metabolic engineering of this versatile chassis organism.

## INTRODUCTION

Biological engineering faces unique challenges that must be addressed before it will be as rapid, reliable, and robust as traditional engineering disciplines (1, 2). On one front, simpler and more flexible genetic engineering tools can facilitate re-writing the genome sequences of living cells to speed up the design-build-test cycle. Another major challenge is the complexity inherent in using living cells as a starting point. Even ‘simple’ microbial cells maintain thousands of distinct components, from small molecules to proteins, but only a smaller subset of these are absolutely required for cellular replication. In fact, many of these functions are unnecessary or even undesirable in controlled laboratory and industrial settings that are not as varied as the natural environments that shaped the evolution of these cells. Therefore, a major aim of the field of synthetic biology has been to re-factor and re-engineer genomes to create ‘chassis’ organisms with reduced complexity (3, 4). The main approaches to genome simplification fall into two categories: a bottom-up approach in which DNA pieces are assembled into a minimal genome that is tested to see if it can successfully ‘boot up’ self-replication in a host cell and a top-down approach in which chromosomal regions found to be dispensable are removed in a step-wise fashion to create a streamlined genome. These efforts have led to more efficient and reliable microbial cell factories and blurred the line between chemical and living systems (5–8).

Bacterial genome streamlining efforts often begin by deleting regions of a chromosome that encode selfish or parasitic sequences that are obviously unnecessary and may even be hazardous to a cell. Deleting transposons and prophage sequences reduces mutation rates and improves the fitness of cells (9–13). It is also relatively straightforward to recognize and then delete gene clusters encoding resource- or energy-intensive functions that cells perform to survive in the wild but are not needed in laboratory or industrial cultures. For example, bacterial growth and productivity has been improved by deleting motility systems, such as flagella (14), and large genomic islands encoding pathways for synthesizing secondary metabolites (15, 16).

For more extreme genome reduction, information about the conservation of genes within a bacterial species and what single-gene knockouts are viable can be integrated to predict additional genome regions to target for deletion (10, 17, 18). Computational tools are emerging that further integrate gene essentiality predictions with whole-cell metabolic and gene expression models to predict the viability of strains that combine multiple deletions (19). Still, viable routes to extreme genome reduction remain difficult to predict because there can be interactions between genes that make some combinations of deletions inviable (e.g., synthetic lethality). Even when multiple large chromosomal regions can be removed, significant growth defects often result from accumulating losses of ‘quasi-essential’ genes, defined as genes that are not needed for survival but are required for robust growth (3, 8, 18, 20, 21). Given how difficult it is to predict how gene essentiality and quasi-essentiality generalizes across environments (22), it is often necessary to empirically test the viability and performance of many different deletions and combinations of those deletions to create a useful minimal genome organism.

*Acinetobacter baylyi* ADP1 is a non-pathogenic soil bacterium that is notable for its naturally transformability under normal laboratory growth conditions (23, 24). ADP1 has been used in fundamental studies of DNA uptake and mechanisms of bacterial genome evolution (25–27). Multiple studies have also highlighted its useful innate metabolic abilities, particularly related to degradation of aromatic compounds (28–32) and production of triacylglycerols and wax esters (33, 34). ADP1’s natural transformability has been leveraged to perform targeted mutagenesis and disruption of genes in its chromosome for metabolic engineering of these pathways (35, 36), as well as to add novel genes to its genome and evolve them to improve these functions (33, 34, 37). The natural transformability of ADP1 makes it an especially tractable system for testing genome streamlining approaches and their impact on engineered functions.

We previously began the process of streamlining the genome of *A. baylyi* ADP1 by creating a transposon-free strain that exhibited lower mutation rates, enhanced transformation, and reduced autolysis (11). Here, we generated a new Tn-Seq dataset defining gene essentiality in rich medium and used this information to design 55 multiple-gene deletions covering 58.3% of its chromosome. To facilitate engineering these large deletions, we developed a “Golden Transformation” genome engineering procedure that uses Golden Gate Assembly (38, 39) to construct the necessary DNA modules for transformation. Nineteen of these large deletions, together comprising 24.6% of the ADP1 genome, were viable. Golden Transformation can also be used to facilitate other genome engineering tasks, such as constructing and inserting gene expression modules into the ADP1 chromosome. Finally, we demonstrate that the native CRISPR/Cas locus in *A. baylyi* ADP1 is active and show that it can be reprogrammed to block reacquisition of deleted genomic regions. Overall, our work demonstrates how the native genetic capabilities of *Acinetobacter baylyi* ADP1 make it an especially tractable system for testing combinatorial routes to more extreme genome streamlining and engineering.

## MATERIALS AND METHODS

### Culture conditions

*Acinetobacter baylyi* ADP1 (30) and ADP1-ISx (11) were grown at 30°C in LB-Miller (10 g NaCl, 10 g tryptone, 5 g yeast extract per liter). Liquid cultures were incubated in 8×150mm test tubes with orbital shaking at 200 r.p.m. over a 1-inch diameter. Solid media contained 1.6% agar (w/v). Media were supplemented with 50 μg/mL kanamycin (Kan), 60 μg/mL spectinomycin (Spec), 20 μg/mL gentamycin (Gent), 200 μg/mL 3-azido-2′,3′-dideoxythymidine (AZT), and 0.3 mM diaminopimelic acid (DAP), as indicated.

### Golden Gate assembly

DNA fragments for transformation were constructed using Golden Gate Assembly (GGA) reactions containing 1 U/µl BsaI-HF or 0.5 U/µl BsmBI and 150 U/µl of either T7 or T4 DNA ligase in T4 DNA ligase buffer (New England Biolabs) (38, 39). The standard ∼5-hour protocol was 31 cycling steps, each consisting of 37°C for 5 min and 16°C for 5 min, followed by one-time final incubations at 55°C for 5 min and 80°C for 5 min. For the short ∼1-hour protocol, the 37°C and 16°C steps were reduced to 1 min apiece. The pBTK622 part plasmid was constructed by adding the *tdk-kanR* cassette (23) to the pYTK001 entry vector as described elsewhere (39). Plasmid samples input into GGA reactions were isolated from *E. coli* cultures using the PureLink Quick Plasmid Miniprep Kit (ThermoFisher). Input PCR products were purified using the GeneJet PCR purification kit (ThermoFisher). Concentrations of all DNA parts were determined using the Qubit dsDNA BR Assay Kit (Invitrogen).

### ADP1-ISx transformation

ADP1-ISx glycerol stocks were streaked out on LB agar and incubated overnight. For each transformation assay replicate, a different colony was picked and inoculated into LB and grown overnight to produce a culture ready for transformation. Transformations in liquid culture were initiated by mixing input DNA, 250 µL LB, and 17.5 µL of the overnight ADP1-ISx culture. Puddle transformations were initiated by mixing input DNA with 50 µL of the overnight ADP1-ISx culture and transferring the complete volume onto a 13 mm diameter, 0.025 µm pore-size mixed cellulose esters membrane (Millipore #VSWP01300) that was left resting on the surface of an LB agar plate. In both cases, transformations were incubated for 24 h under normal *A. baylyi* growth conditions. Then, dilutions in sterile saline were plated on LB and LB-Kan agar to estimate transformation frequencies or on LB-Kan or LB-AZT agar for genome modification steps.

### Transformation frequency assays

Overlap extension PCR (OE-PCR) transformations used 0.01 pmol of a 3,875-bp DNA product constructed by joining the *tdk*-*kanR* cassette to 1.2 kb of 5’ homology and 1.0 kb of 3’ homology to Site 2 in the ADP1 genome (11). GGA DNA transformations used 10 µL assembly reactions that included 0.01 pmol of each DNA part: the pBTK622 plasmid and PCR amplicons that added the necessary restriction sites to the same two flanking genomic homology regions. Multiple GGA reactions were pooled and re-divided when testing the same DNA construct in multiple replicates and across conditions. Primer sequences for creating the PCR products used in each type of assembly are provided in **Table S1**.

Transformation frequencies were determined by comparing colony-forming units (CFUs) on LB-Kan agar to CFUs on LB agar with 3-4 replicates for each condition. Puddle transformations were resuspended by carefully transferring the filter with sterile tweezers into 10 mL of saline and vortexing to release cells before plating dilutions. Negative controls without DNA yielded no colonies on LB-Kan agar. Transformation frequencies and confidence intervals were estimated from CFU values using Poisson regression models in R (40).

### Tn-Seq

The transposon mutagenized library of *A. baylyi* ADP1 was constructed through conjugation of the mini-transposon suicide vector pBT20 (41). For the conjugation, overnight cultures of *E. coli* β2163 (42) carrying pBT20 and ADP1 were pelleted via centrifugation and resuspended together at a 3:1 ratio in 100 µL of sterile saline. This mixture was then spotted onto the same type of membrane used in puddle transformations and placed on an LB-DAP plate. After ∼24 hours at 30°C, the filter was transferred to a 1.7 mL Eppendorf tube with 1 mL of sterile saline. After vortexing to resuspend cells, 800 μL of this mixture was used to inoculate 50 mL of LB-Gent. After overnight growth, 1.2 mL aliquots of this culture were stored at −80°C as glycerol stocks.

For sequencing library preparation, genomic DNA was extracted from the frozen aliquots using a Purelink Genomic DNA Mini Kit (ThermoFisher) and sheared on a Covaris S220 to an average size of 300 bp. Poly-C tails were added to the fragmented DNA using terminal deoxynucleotidyl transferase (Promega). Then, the processed fragments were size-selected using AMPure XP beads (Beckman Coulter) and amplified by PCR with a biotinylated primer binding inside of the transposon and a non-biotinylated primer binding to the poly-C tail. The amplified library fragments were then bound to streptavidin M-280 Dynabeads (ThermoFisher) and washed to remove unbound DNA. Illumina barcodes and adaptors were added in a subsequent PCR reaction in which internal barcodes were introduced to allow multiple libraries to be run with the same external Illumina barcode. The libraries were pooled in equal ratios, analyzed on a BioAnalyzer 2100 (Agilent) for fragment distribution, and sequenced on an Illumina HiSeq 4000 at the University of Texas at Austin Genome Analysis and Sequencing Facility (GSAF). Tn-Seq FASTQ files are available from the NCBI Sequence Read Archive (PRJNA559727).

Tn-Seq reads were analyzed as previously described (43). Briefly, reads containing transposon sequences were quality filtered, trimmed of adapter sequences, and then mapped to the *A. baylyi* ADP1 genome (GenBank: NC_005966.1) (30). To identify essential genes, we compared the observed number of insertion mutants in each gene to simulated null distributions based on the total number of possible insertion sites. Genes designated as essential were significantly depleted for transposon mutants when compared to the simulated null distributions using DESeq2 (44). Scripts for performing this analysis are freely available online (https://github.com/barricklab/Tn-seq). Genes in the *A. baylyi* ADP1 genome were assigned predicted functional categories using eggNOG-mapper (version 4.5.1) (45). The full output of the Tn-Seq analysis with COG assignments for each gene is shown in **Table S2**.

### Multiple-Gene Deletion Strain Construction

Large deletion variants of ADP1-ISx were generated by transforming GGA reactions, with the exception of MGD7 and MGD11, which were generated by transforming OE-PCR reactions. GGA reactions and transformations were performed as described above except we used 40 µL GGA reactions containing 500 ng of pBTK622 and 600 ng of each of the ∼2-kb flanking genomic homology fragments generated by PCR. When standard transformations of a GGA reaction for a deletion yielded no colonies or very few colonies on LB-Kan agar, we attempted puddle transformation of a new GGA reaction.

### Genome Sequencing

Genomic DNA isolated from candidate large-scale deletion strains was prepared for sequencing as described previously (46). Briefly, genomic DNA was extracted from overnight cultures, then fragmented, end-repaired, A-tailed, ligated to adapters, and size-selected for library preparation. The resulting DNA libraries were sequenced to >100× coverage on an Illumina HiSeq 2500 platform at the UT Austin GSAF. Read files were analyzed with *breseq* (version 0.28.0) (47, 48) to predict point mutations, large deletions, and other types of structural variation relative to the *A. baylyi* ADP1 genome. FASTQ files from genome sequencing are available from the NCBI Sequence Read Archive (PRJNA559727).

### Growth rates

The growth rates of multiple-gene-deletion strains that still contained the *tdk*-*kanR* cassette in their genomes were compared to growth rates of ADP1-ISx. For each strain being tested, 2 µL of a glycerol stock was inoculated into 200 µL of LB in a 96-well clear flat-bottom microplate (Costar). This microplate was incubated overnight at 30°C with 250 r.p.m. orbital shaking over a diameter of 1-inch. Subsequently, 2 µL of the preconditioned well for each strain was used to inoculate 200 µL of fresh LB in each of three replicate wells in a new microplate. The optical density at 600 nm (OD600) was recorded every 10 minutes following a brief shaking step (15 seconds with a 6 mm amplitude) as these cultures were incubated at 30°C in an Infinite M200 Pro microplate reader (Tecan).

OD600 values were corrected by subtracting measurements of blank wells and then further normalized by offsetting all measurements in each well such that the first three points had the same average as the grand mean of these points across all inoculated wells on the entire plate. Growth rates were determined from nonlinear least squares fits to an exponential growth equation with lag and doubling time parameters in R (40). The maximum specific growth rate fit from any set of 7 consecutive points (spanning 1 h of measurements) that all had OD600 values between 0.01 and 0.15 was determined for each well.

### CRISPR assays

The ‘CRISPR-Ready’ variant of ADP1-ISx was created by replacing all but one of the 91 spacers of its native CRISPR array with the *tdk*-*kanR* cassette using BsaI Golden Transformation and selection on LB-Kan. CRISPR reprogramming was accomplished by transforming the CRISPR-Ready strain with a rescue cassette encoding a new spacer sequence. To assemble each spacer cassette, two complementary oligonucleotides were synthesized and annealed to create a double-stranded insert flanked by restriction sites. The spacer was then added to the genome via BsmBI Golden Transformation using the same two homology flanks used in the replacement step and selection on LB-AZT agar. Proper insertion of each spacer into the ADP1-ISx genome was verified by PCR and Sanger sequencing. Transformation assays for acquisition of a spectinomycin resistance gene were conducted as described above but using 100 ng of donor DNA from an ADP1 strain constructed by replacing bases 139832–140099 of the genome (GenBank: NC_005966.1), which overlap the nonessential gene *ACIAD0135*, by transforming an OE-PCR product containing the *specR* cassette from pIM1463 and flanking homology (49).

In the CRISPR-Lock experiment, CRISPR-Ready variants of the MGD2 and MGD18 deletion strains and an ADP1-ISx strain with a suspected genome rearrangement were constructed as described above. Self-targeting spacer cassettes were created by BsmBI GGA as described above. AZT-resistant colonies were further characterized for mutations in the *tdk*-*kanR* cassette versus spacer acquisition by patching to determine if they remained Kan resistant and by PCR and Sanger sequencing, if necessary. Oligonucleotides used to conduct these experiments are provided in **Table S1**.

## RESULTS

### Golden Transformation

*A. baylyi* ADP1 is highly competent for DNA uptake during normal growth. Transformation frequencies of >10^−2^ per cell can be achieved for genomic DNA from a donor ADP1 strain with a genetic marker embedded in its chromosome (24), but transformation is less efficient for DNA fragments with shorter flanking homologies. Successful integration of a foreign DNA sequence into the genome, which occurs most readily via RecA-mediated recombination, is generally the limiting step for transformation (50). Therefore, robust protocols for engineering the genome of ADP1 typically require assembling linear double-stranded DNA cassettes with at least 500 base pairs of flanking genomic homology added on each end, and using even longer flanking homology regions further increases the efficiency of transformation (50, 51).

Currently, the most common approach used for ADP1 genome engineering relies on PCR assembly of DNA constructs for transformation. This procedure has variously been called overlap extension PCR (OE-PCR), splicing PCR, fusion PCR, or other names (23, 31). In this method, one first generates a marked replacement DNA cassette containing dual positive and negative selection functionalities (e.g., *tdk* and *kanR* genes) flanked by DNA sequences matching the genomic target site. The PCR product is added to a growing culture of ADP1, allowing DNA uptake and homologous recombination to replace the native sequence between the homology regions. Successful transformants are isolated by plating on agar containing the positive selective agent (e.g., kanamycin). In a second step, an unmarked rescue cassette containing flanking homologies to the insertion site is added to accomplish another recombination event with the chromosome. The rescue cassette can be designed to precisely remove the replacement cassette and create a deletion or to insert new genes into the genome at that site. Successful rescue transformants are isolated by plating on agar containing the negative selection agent (e.g., AZT).

OE-PCR uses multiple PCR reactions with overlapping primers designed to stitch together longer pieces of DNA. In our experience, this is an unreliable and time-consuming method for creating the DNA cassettes needed for ADP1 genome editing. OE-PCR reactions may yield off-target amplicons and then require optimization of PCR conditions or gel purification of intermediate PCR fragments to produce a clean DNA assembly, which becomes cumbersome for large projects. To streamline this process, we tested using Golden Gate Assembly (GGA) to create the replacement and rescue cassettes needed for *A. baylyi* genome engineering more efficiently and quickly. Fundamentally, GGA uses plasmids or PCR fragments encoding parts that are flanked by recognition sites for a type IIS restriction enzyme (38, 39). The restriction enzyme cut sites are positioned to generate compatible overhangs between fragments. Multiple DNA pieces can then be assembled in a designed order in a one-pot reaction that contains both a restriction enzyme and DNA ligase.

For our method, which we dubbed Golden Transformation (GT), we use GGA to combine one or more DNA parts with the genome homology regions required for *A. baylyi* genome engineering (**Fig. 1**). In our design, just two PCR reactions that amplify the flanking ∼1-kb homology regions from genomic DNA are needed. Each PCR contains a primer that adds terminal BsaI and a BsmBI restriction sites. For the first “replacement” step of genome modification, these flanking regions are combined with the *tdk*-*kanR* cassette from plasmid pBTK622 in a BsaI GGA reaction. This DNA product is used to transform ADP1 to insert the *tdk*-*kanR* cassette in place of any genes that are to be deleted. For the second “rescue” step, the same two flanking PCR fragments amplified from the genome are used in a GGA reaction. If a deletion is desired, the two PCR products can be joined together with only a 4-bp scar remaining in a BsmBI GGA reaction. Alternatively, since the BsaI and BsmBI overhangs can be designed to be compatible with GGA-based toolkits of genetic parts (39, 52), one can create and insert a transcriptional unit or a larger construct into the genome in place of the *tdk*-*kanR* cassette.

**Figure 1.**
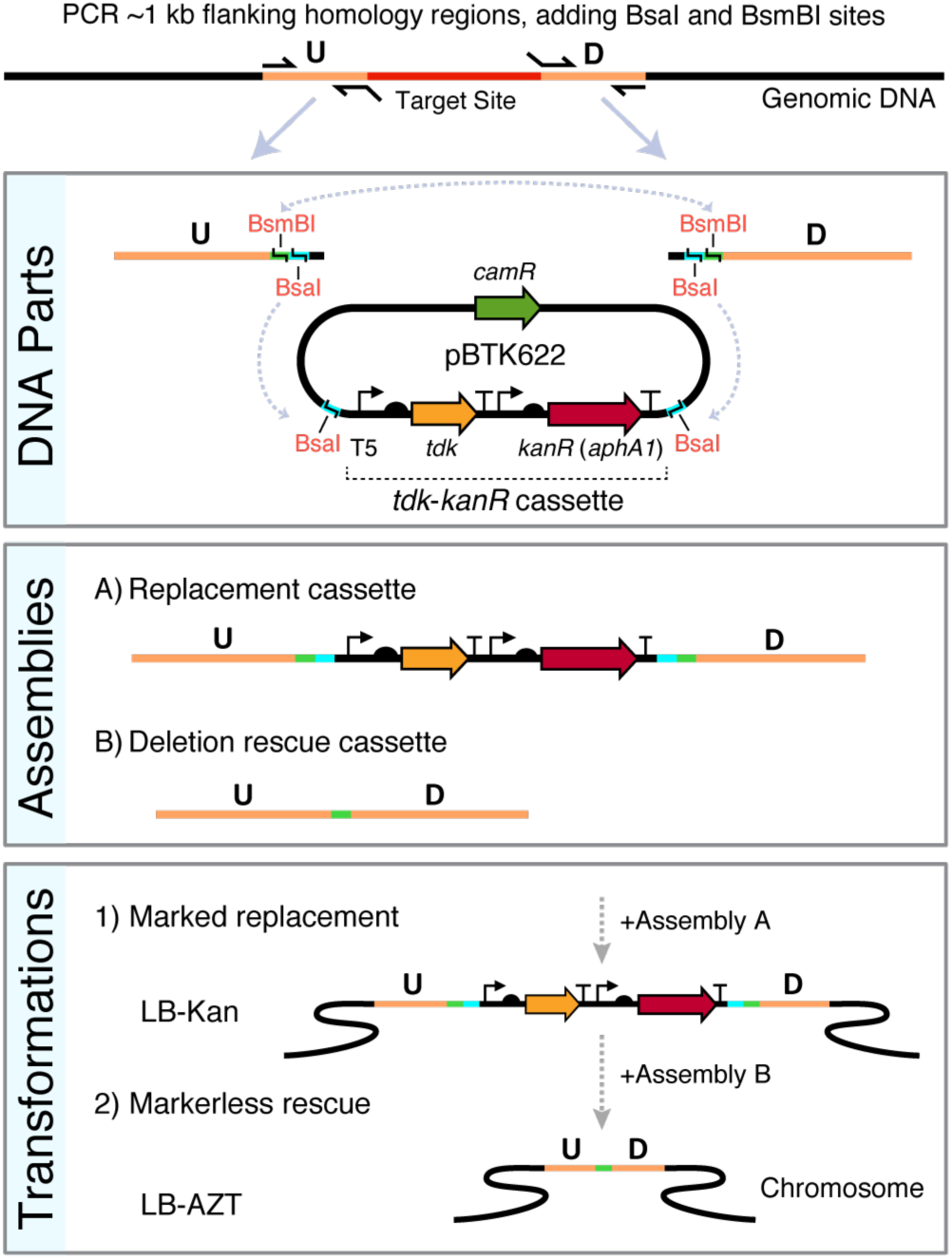
Golden Transformation method for ADP1 genome engineering. Two PCR reactions are performed to create upstream (U) and downstream (D) genomic target flanks with added terminal BsaI and BsmBI type IIS restriction sites as depicted. The two PCR products can then be combined via BsaI Golden Gate assembly (GGA) with the selection cassette to form a replacement DNA or combined with one another and optionally with additional genetic parts (not shown) via BsmBI GGA to form a rescue cassette. The positive-negative selection cassette (*tdk*-*kanR*) is maintained on the high-copy pBTK622 plasmid that has an origin that does not replicate in *A. baylyi*. The first GGA reaction is added to an *A. baylyi* culture and then plated on LB-Kan to select for transformants with the replacement cassette integrated into the genome. Then, transformation of the second assembly reaction with counterselection on LB-AZT is used to move the unmarked deletions/additions encoded on the rescue cassette into the genome.

### Testing Golden Transformation

We first tested the efficiency of GT using “replacement” DNA assemblies that included the *tdk*-*kanR* cassette and flanking genome homology regions (**Fig. 2**). OE-PCR product was used as a benchmark for the expected maximum efficiency that would be possible with the GT procedure. It yielded transformation frequencies of 1.7 × 10^−4^ per cell, which were similar to previous tests of this construct (11). Adding GGA buffer containing heat-inactivated T7 ligase and BsaI to the OE-PCR product only slightly reduced transformation: by a factor of 1.8-fold [1.3–2.3] (95% confidence interval). Therefore, we concluded that it was not necessary to purify GGA reactions before they were added to cells for transformation.

**Figure 2.**
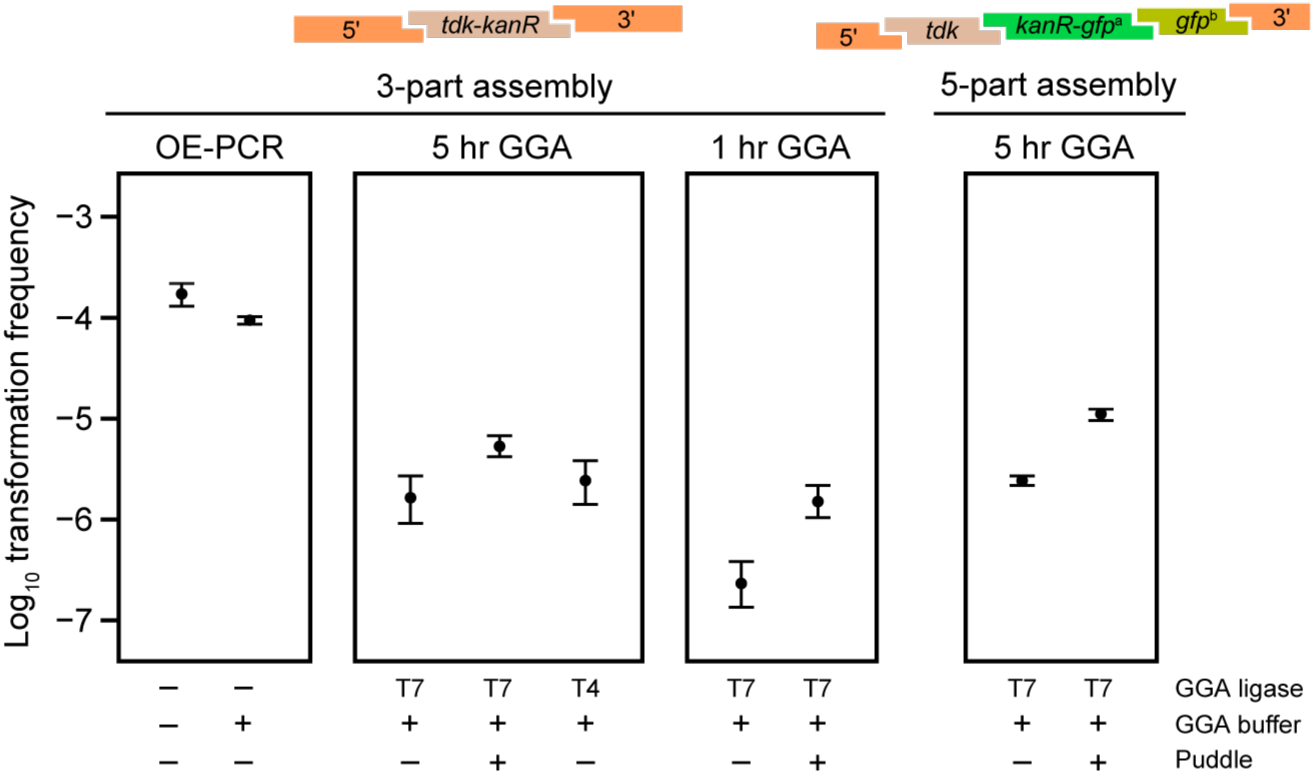
Golden Transformation can achieve high genome editing rates. Transformation of two different cassettes constructed in 3-part or 5-part Golden Gate assembly (GGA) reactions were compared by counting colonies obtained on LB-Kan versus LB agar. Transformations of a purified DNA sample constructed by overlap-extension PCR (OE-PCR) at a concentration that corresponds to 100% efficient assembly of the 3-part GGA reaction and this OE-PCR sample with GGA buffer added to it were included for comparison. The effects of “short” (1 hr) or “long” (5 hr) GGA thermocycling programs, using T4 versus T7 ligase in GGA reactions, and performing “puddle” transformations that concentrate cells and DNA by combining them on a filter placed on an agar surface rather than in a liquid culture were also tested.

Golden Transformation using standard 5-hour GGA reactions with three input DNA pieces yielded transformation frequencies that were reduced by roughly two orders of magnitude relative to the OE-PCR benchmark representing the expectation for 100% efficient DNA assembly. Due to the high transformability of *A. baylyi* ADP1, this still represents thousands of transformants per standard GT assay. There was very little difference in transformation frequencies whether T4 or T7 ligase was used in GGA. Shortening the steps in the GGA temperature cycling protocol to a shorter 1-hour program reduced the transformation frequency by a factor of 6.9, consistent with DNA assembly limiting GT efficiency. Interestingly, GT of a larger five-part assembly was roughly as efficient as GT of the three-part assembly. The ability to create this type of complex cassette and integrate it into the *A. baylyi* genome in a single step demonstrates a further advantage of the GT strategy over using OE-PCR for DNA assembly.

We next tested whether one could compensate for the reduced efficiency of GT by performing “puddle” transformations. In this setup, *A. baylyi* and DNA are combined on a filter that is transferred to an agar surface for cell growth, as opposed to the normal transformation procedure in which DNA is added to cells in a liquid culture. The puddle procedure improved the transformation frequency for each of three GGA reactions that were tested by a similar amount. The group-wise increase by a factor of 5.3 [4.1–6.8] (95% confidence interval) matched the 5-fold higher amount of DNA per cell in puddle transformations compared to the standard procedure. The absolute yield of successfully transformed cells is also higher in puddle transformations, despite a reduction in the total number of cells present after growth by a factor of 1.9, on average, compared to the transformation assays in liquid culture.

### Tn-Seq to Determine ADP1 Gene Essentiality in LB

To demonstrate how Golden Transformation can be used to rapidly construct deletions, we applied it to an *A. baylyi* ADP1 genome streamlining project. The ADP1 genome sequence is annotated with 3305 protein-coding genes and 100 RNA genes (30). Previously, a collection of single-gene deletion strains was constructed by replacing individual open reading frames with the *tdk*-*kanR* cassette (53). This study was able to classify 499 proteins as essential and 2593 as dispensable for ADP1 viability in a minimal succinate growth medium. We used transposon sequencing (Tn-Seq) to determine which ADP1 genes are required for robust growth in rich medium (LB), so that we could identify regions of the genome that were candidates for large deletions that would remove many genes at once while still preserving growth under a broad range of conditions.

Sequencing >4 million reads in the Tn-Seq library, corresponding to 59,757 unique insertion sites, enabled us to confidently classify 2871 proteins as either essential or dispensable to ADP1 fitness in LB (**Fig. 3A**, **Table S2**). Comparing this set to the knockout collection, 238 proteins were classified as essential in both experiments, 261 were essential for viability in minimal medium only, 108 were essential for fitness in rich medium only, and 2231 were dispensable in both experiments. Assigning ADP1 proteins to Clusters of Orthologous Groups (COGs) showed that certain functions were overrepresented within each of these categories (**Fig. 3B**). As expected, proteins with translation, ribosomal structure, and biogenesis functions (J) are most common in the shared essential group, and proteins in the shared dispensable group commonly have an unknown function (S) or are not even assigned to a COG (X). Amino acid transport and metabolism functions (E) predominate among those proteins that are essential for growth in minimal medium and dispensable for maintaining fitness in rich medium, presumably because many biosynthetic pathways are unnecessary when amino acids are supplied as nutrients. Genes essential for fitness in LB yet dispensable in minimal succinate medium are largely involved in energy production and conversion (C). This observation is consistent with ADP1 needing to utilize a more diverse pool of compounds for energy generation in rich medium.

**Figure 3.**
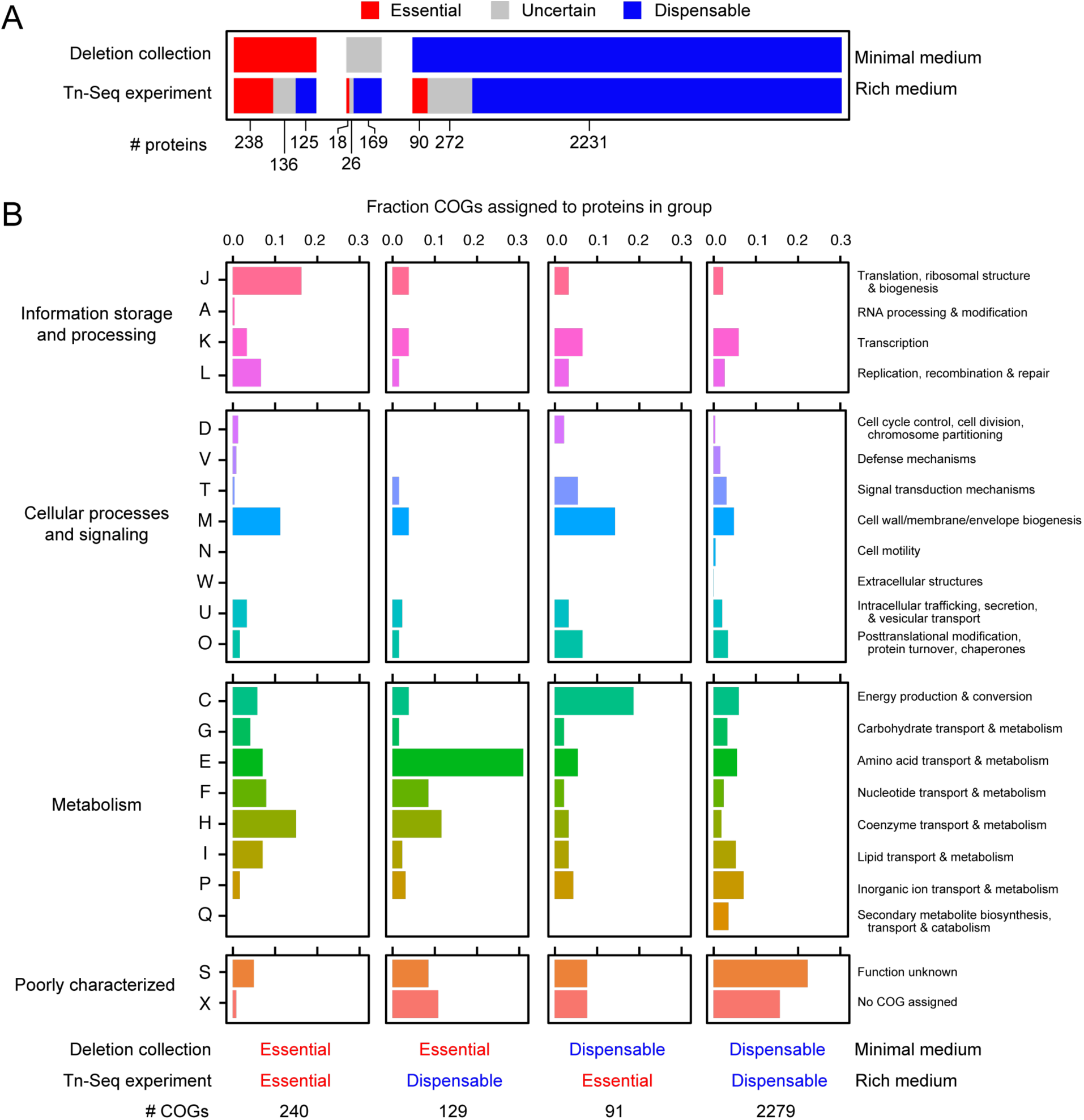
*A. baylyi* ADP1 protein essentiality. (A) Comparison of the essentiality of protein-coding genes between two experiments. The deletion collection determined which strains lacking a single protein remained viable in a defined minimal succinate medium (minimal medium) (53). The Tn-Seq experiment from the current study examined the representation of transposon insertions in different genes within populations of mutants cultured in LB (rich medium), which reveals proteins that are essential for fitness in this environment. Colored bars correspond to the numbers of proteins that were judged as essential, dispensable, or uncertain (meaning either untested or ambiguous). The horizontal bars are broken up to show the overlap of proteins classified into each category between experiments. (B) Functions of proteins classified as various combinations of essential or dispensable across the two experiments. Each column shows the breakdown of Clusters of Orthologous Groups (COG) functions predicted for genes that were definitively classified in each experiment (not in the uncertain category).

### Multiple-gene-deletion Strains

We used information about gene essentiality from the knockout collection and our Tn-seq results to design 55 multiple-gene deletions covering a total of 2.13 Mb (59.4%) of the 3.59 Mb ADP1-ISx genome (**Fig. 4A**, **Table S3**). In addition to spanning as many nonessential genes as possible, some of the regions targeted for deletion included one or a few genes that were predicted to be essential in one or both growth conditions. We expected that some of these genes might be conditionally dispensable (i.e., able to be deleted if other nearby genes involved in the same process were removed at the same time). For example, the DNA methylase in a restriction-modification system is essential for cell viability unless its corresponding restriction enzyme is also deleted. Similar interactions can arise in metabolic pathways that have toxic intermediates and in other coupled cellular processes.

**Figure 4.**
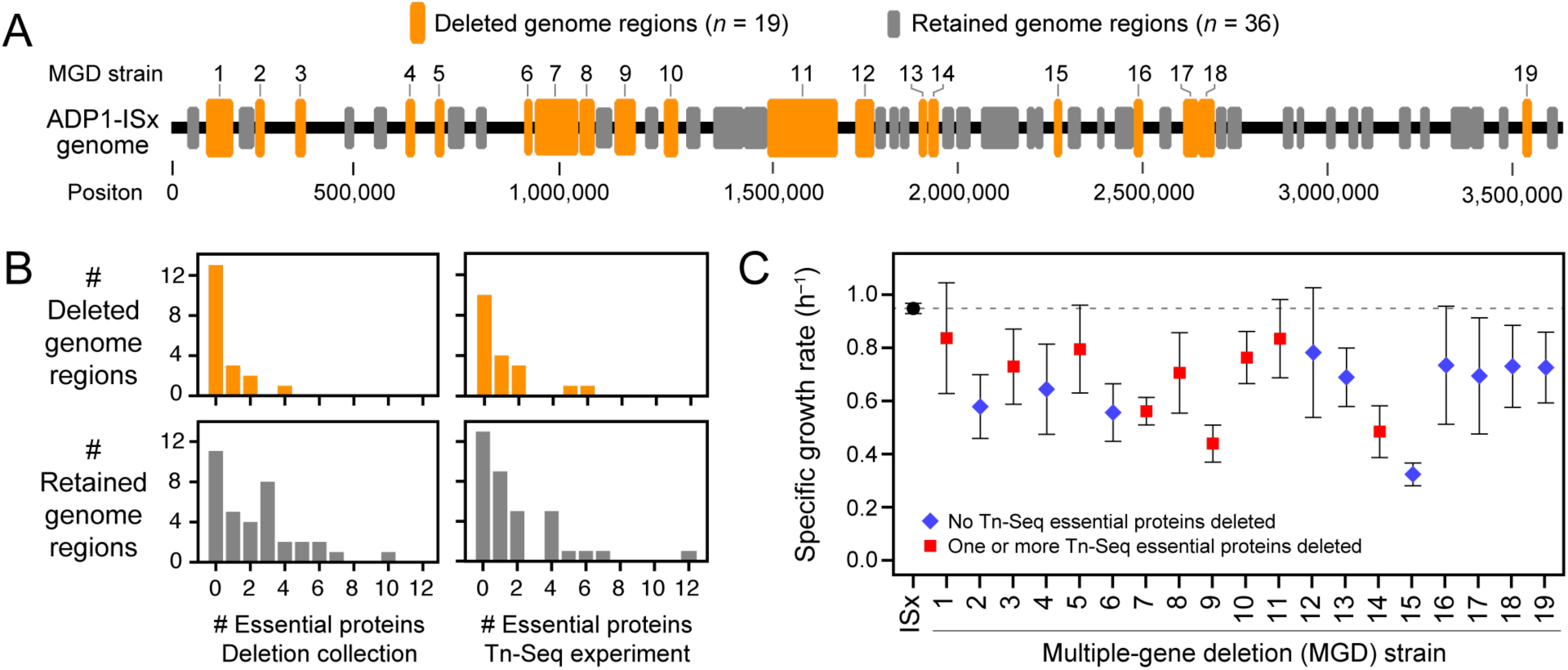
Dispensability of *A. baylyi* genome regions targeted for deletion and growth rates of multiple-gene deletion strains. (A) Regions targeted for deletion from the ADP1 chromosome. Successful deletions that resulted in multiple-gene deletion (MGD) strains are numbered and displayed in orange. Deletions that could not be constructed are shown in gray. (B) Gene essentiality in deleted and retained regions. Histograms show the breakdown of the 19 deleted regions and the 36 retained regions by how many proteins they included that were classified as essential in the deletion collection or the Tn-Seq experiment. (C) Maximum specific growth rates of MGD strains in LB determined from fitting growth curves monitored by optical density in a microplate reader. Shapes and colors of symbols indicate whether any essential proteins from the Tn-Seq experiment were included in that deletion. Error bars are 95% confidence intervals.

We attempted construction of the 55 planned ADP1-ISx derivatives with multiple-gene-deletions using Golden Transformation in rich medium (LB). Single-gene deletion studies commonly encounter problems with false-positives, such as integration of the selectable marker into one copy of a region of the genome that has been transiently amplified in the target cell (54–56). To detect these artifacts, we validated that the *tdk*-*kanR* selection cassette inserted into the genome at the expected site and replaced the targeted genome region in candidate deletion strains by performing several PCR reactions and, in some cases, by re-sequencing whole genomes (**Fig. S1**, **Table S3**). In total, 19 multiple-gene-deletion (MGD) strains were successfully created. Collectively, they dispense with 24.6% of the ADP1-ISx genome.

For 10 of the 36 regions that we could not delete—which we designate as “retained genome regions” (RGRs)—we did not obtain any transformants following GT, and there were generally 10- to 100-fold fewer transformants in GTs for the other 26 RGRs compared to those that yielded successful deletions (**Table S3**). Of these 26 putative deletions that subsequently failed PCR verification, most appeared to have either one-sided integrations of the *tdk*-*kanR* cassette, with rearrangements or incomplete deletions on the opposite side, or to have large chromosomal amplifications which enabled one copy of the targeted genes to be replaced by the *tdk*-*kanR* cassette while leaving one or more other copies intact. The *A. baylyi* genome is known to experience gene amplifications at high rates (25, 57), which led to false-positives during construction of the single-gene knockout collection (53). To check for these or other types of false-positives in the MGD strains, we validated a subset via whole-genome sequencing, including as controls two strains that failed the PCR assays. All 12 of the sequenced MGD strains were found to have the expected deletions. Genome sequencing corroborated that the other two strains, which had been transformed with DNA targeting either RGR16 or RGR28 for deletion, had one-sided homologous integrations of the *tdk*-*kanR* cassette. In each case there was a deletion of only a few hundred bases on opposite (nonhomologous) side of the cassette.

We further examined whether the presence of one or more proteins found to be essential in the minimal-medium single-gene knockout collection or in the rich-medium Tn-Seq experiment was predictive of whether a deletion was ultimately successful (**Fig. 4B**). Deleted genome regions were significantly less likely to contain at least one gene flagged as essential in the single-gene knockout collection as regions that we were unable to remove (*p* = 0.008, respectively, one-tailed Fisher’s exact test). There was also a lower chance of having a gene designated essential by the Tn-Seq experiment within a successfully deleted region, but not significantly so (*p* = 0.186). The worse correlation with the Tn-Seq experiment most likely reflects that essentiality, as measured by this assay, is for retaining high fitness rather than absolute viability for growth.

### Growth characteristics of MGD strains

We next tested the viability and growth rates of the deletion strains. They were constructed in LB, so they all can grow in this rich medium. However, nearly all deletion strains had moderate 10-25% reductions in maximum specific growth rate in LB compared to the ancestral ADP1-ISx strain, and the MGD9, MGD14 and MGD15 strains exhibited larger reductions of 50% or more (**Fig. 4C**). The magnitudes of these growth defects were not predictable from the overall characteristics of the deletion. For example, there was not a significantly different growth rate depending on whether a deletion contained a protein classified as essential in the Tn-Seq experiment or in the deletion collection (*p* = 0.29 and *p* = 0.40, respectively, two-tailed *t*-tests). There was also no consistent correction between the size of the genome region deleted and the growth rate of a strain (*p* = 0.31, for a non-zero slope when bootstrap resampling by strain). Finally, only 2 of the 19 deletion strains (MGD9 and MGD15) showed no growth when cultured in minimal succinate medium for 72 hours.

### Reprogramming the native CRISPR-Cas system of *A. baylyi* ADP1

The ADP1 genome encodes a complete type I-Fa CRISPR-Cas system (58). We tested whether this system was functional by reprogramming it to target a foreign DNA sequence (**Fig. 4A**). First, we replaced the entire native CRISPR array of spacers that is proximal to the Cas-operon with the *tdk*-*kanR* cassette (**Fig. 5A**). We refer to an *A. baylyi* strain with this modification as “CRISPR-Ready”. One or more custom spacers can be re-inserted to reprogram the CRISPR-Ready strain with a transformation that uses AZT counterselection against *tdk* (**Fig. 5B**).

**Figure 5.**
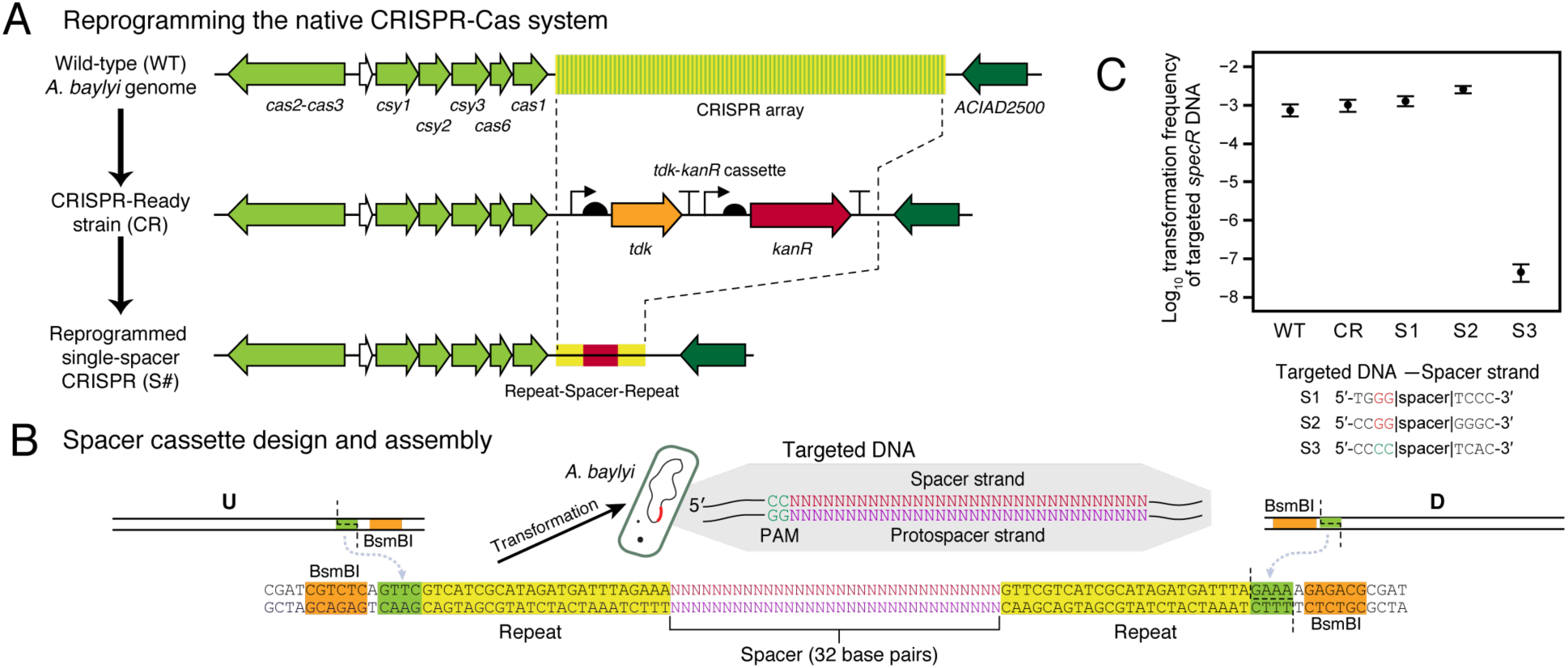
The native *A. baylyi* ADP1 CRISPR-Cas system is active and can be retargeted.

A. Scheme for reprogramming the *A. baylyi* CRISPR array. First, a “CRISPR-Ready” strain is created by performing a Golden Transformation that replaces the entire native spacer array with a *tdk*-*kanR* cassette. Then, a second Golden Transformation can be used to add a rescue cassette that contains one or more designed spacers under control of the native gene expression signals.
B. Single-spacer replacement cassette design. Synthetic double-stranded DNA encoding the spacer and surrounding repeats is combined with PCR products corresponding to the flanking genome homology upstream (U) and downstream (D) using BsmBI Golden Gate assembly. The inset shows the DNA sequence that is targeted for cleavage with the protospacer adjacent motif (PAM) typical of type-I CRISPR-Cas systems. (C) Reprogrammed CRISPR-Cas system restricts transformation of foreign DNA. Frequencies of transformants of genomic DNA from *A. baylyi* ADP1 donor that has an integrated spectinomycin resistance gene (*specR*) were used to judge whether targeting a spacer to this sequence in a recipient strain prevented its acquisition. WT is wild type ADP1-ISx and CR is the CRISPR-Ready derivative of this strain. S1 and S2 are controls with spacers that match the *specR* sequence but in incorrect PAM contexts. S3 is an on-target *specR* spacer with the correct PAM. Error bars are estimated 95% confidence intervals.

The protospacer adjacent motif (PAM) for type I CRISPR-Cas systems has been reported to be a downstream GG dinucleotide (59), including for the subtype I-F system of *Acinetobacter baumanii* (60). This is equivalent to a CC located 5’ of the spacer sequence on the strand that it matches in the target site DNA sequence. We found that matches for some of the native *A. baylyi* spacers to *Acinetobacter*-related phages supported this PAM sequence and orientation using the CRISPRTarget software (61). Therefore, we tested the ability of an *A. baylyi* ADP1-ISx strain reprogrammed with a single spacer matching a site in a spectinomycin-resistance gene sequence (*specR*) with this PAM to prevent transformation of this genetic marker when DNA isolated from a strain with the *specR* gene integrated into its genome was added to cultures (**Fig. 5C**). As controls, we tested wild-type, the CRISPR-Ready derivative of this strain, and also two variants in which the CRISPR array was re-targeted to *specR* gene sequences that had incorrect PAM sequences. The properly re-targeted strain reduced transformation frequency by a factor of 10^5^, whereas the transformation frequency was unchanged in the other strains, indicating that the native *A. baylyi* CRISPR-Cas system is functional and highly efficient.

### Assuring and securing deletions with a CRISPR-Lock

We next considered how CRISPR spacers targeting the chromosomal regions removed in a multiple-gene deletion strain could be used for two purposes to further the ADP1 genome streamlining project. First, we tested whether it was possible to add a deletion-targeting spacer to the genome for assurance: to verify whether a gene has been successfully eliminated from a candidate MGD strain. Second, if one has two MGD strains, one with a spacer targeting its own deletion and the other that still retains the *tdk*-*kanR* cassette replacing its deletion, then genomic DNA from the latter can be used to transform the former to readily combine the deletions in one strain. In this case, the deletion-targeting spacer serves as a “CRISPR-Lock” that prevents restoration of the genes originally deleted in the first strain from the genomic DNA of the second strain. Regaining genes in this way is expected to be strongly selected for when a deletion causes a growth defect.

We tested the assurance scheme by attempting to reprogram CRISPR-Ready strain derivatives with self-targeting spacers (T2 and T18) aimed at sequences that were removed in two of the multiple-gene deletion strains, MGD2 and MGD18, respectively, or with the first spacer from the native CRISPR array (N1) as an off-target control (**Fig. 6**). We would expect to not be able to transform a strain that does not have the matching deletion with a self-targeting spacer because this will lead to cleavage of its own genome and cell death. However, escape mutants can evolve that have spontaneous mutations inactivating the *tdk* gene that give them resistance to AZT without replacement of the *tdk*-*kanR* cassette in the step in which we add the spacer to a CRISPR-Ready strain. There is also the possibility of mutations or errors in the engineered spacer such that it no longer self-targets. The relatively high frequency of ∼10^−6^ of background AZT-resistant (AZT^R^) escape mutants makes it difficult to determine whether the self-targeted region was present in the genome based on measuring differences in transformation frequencies of the different spacers alone. Therefore, we screened 10 colonies for whether they maintained kanamycin resistance (Kan^R^) as a quick proxy for whether these putative spacer replacements were actually escape mutants of this variety.

**Figure 6.**
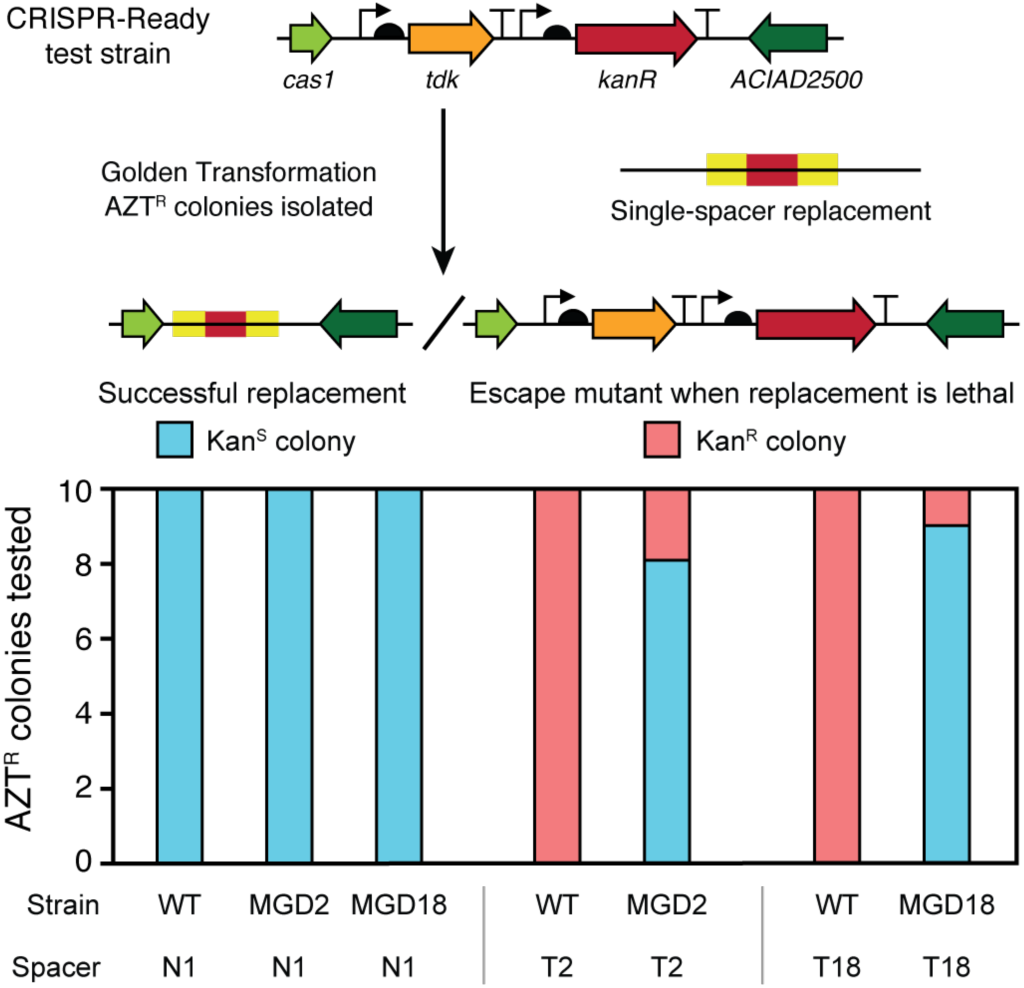
Self-targeting spacers can be used to assure deletions and create a CRISPR-Lock. CRISPR-Ready variants of wild-type ADP1-ISx (WT) and two multiple gene deletions strains (MGD2 and MGD18) were transformed with different spacers to assess the presence of a sequence located within the putatively deleted region. N1 added back the first spacer sequence from the native CRISPR array. It serves as a control because it does not target any sequence in the *A. baylyi* genome. T2 and T18 are spacers that match sequences in the ADP1-ISx genome that are within the regions deleted in the corresponding MGD strains. For each strain-spacer combination tested, 10 AZT^R^ colonies were isolated after transformation with the single-spacer replacement DNA. Successful integration of the spacer can only occur if the targeted region is not present in the recipient strain’s genome. It results in these AZT^R^ colonies also becoming Kan^S^. If integration of the spacer is lethal, then AZT^R^ colonies are expected to have mutations that inactivate the *tdk* gene and remain Kan^R^, as illustrated in the upper panel. Strains with successful spacer integrations from the MGD2+T2 and MGD18+T18 transformations have a “CRISPR-Lock” in their genomes that can prevent re-acquisition of the deleted regions when combining multiple deletions in further stages of the genome streamlining project.

CRISPR-Ready variants of wild-type ADP1-ISx, MGD2, and MGD18 were constructed as detailed above (**Fig. 5**), after first removing the *tdk*-*kanR* cassette from the deleted regions in the latter two strains using GT with the flanking PCR homology segments. Of the AZT^R^ colonies isolated after transformation of each of these strains with the control N1 spacer construct, 10/10 were kanamycin sensitive (Kan^S^) in all three cases, indicating 100% successful integration of this off-target spacer (**Fig. 6**). When we transformed wild-type ADP1-ISx with either of the self-targeting spacers (T2 or T18) all 10 AZT^R^ colonies characterized were now Kan^R^, which the expected result because these spacers should be lethal in the absence of their cognate deletions. In the case of the T18 construct, we found that all 10 colonies had a 551-bp deletion inactivating the *tdk* gene. Most AZT^R^ colonies isolated after transformation of MGD2 and MGD18 with spacers targeting the regions deleted in each strain were Kan^S^ (8/10 and 9/10, respectively). Sequencing confirmed that these isolates had proper spacer integration. The one exception for MGD18 contained a 1-bp deletion in the *tdk* gene. It is also possible for AZT^R^ colonies to be Kan^S^ if they incorporate a mutated copy of the spacer that no longer matches the genome, but we did not observe that outcome. In summary, there were significant enough differences in the chances of getting AZT^R^ colonies that are sensitive or resistant to kanamycin (*p* = 2.6 × 10^−8^, Fisher’s exact test of combined T2 and T18 results) that this assay could be used to diagnose whether a gene was successfully deleted from the *A. baylyi* genome for assurance.

Successful MGD2+T2 and MGD18+T18 transformants from this experiment were used to test the CRISPR-Lock approach. We transformed these strains with genomic DNA from the MGD17 strain still containing the *tdk*-*kanR* replacement cassette. The MGD17 deletion is separated by less than 900 bp from the one in MGD18 (**Table S3**). In each case, 3/3 *kanR* colonies screened had successfully added the new deletion and retained the original one, as determined by PCR and Sanger sequencing. However, we also were able to create the same double deletions by transforming MGD2 and MGD18 strains that did not have the self-targeting T2 or T18 spacers with the same efficiency of 3/3 *kanR* colonies screened having both deletions. Though it was not necessary to block re-incorporation of the deleted segments with the CRISPR-Lock when making these double deletion strains, we expect that it will be increasingly important to prevent re-acquisition of deleted regions as more and more deletions are combined and result in strains with progressively worse fitness as this genome streamlining process continues.

## DISCUSSION

We developed the Golden Transformation procedure to address limitations in the current methods used to assemble DNA cassettes for modifying the genome of the naturally competent bacterium *A. baylyi* ADP1. The ease of Golden Gate assembly combined with the very high transformability of unpurified DNA from these reactions into *A. baylyi* makes it possible to use a procedure that requires fewer steps and fewer oligonucleotides than PCR assembly methods. We demonstrated the utility of Golden Transformation by generating a collection of 19 derivatives of the transposon-free *A. baylyi* ADP1-ISx strain that each have one additional very large chromosomal deletion spanning 20,920 to 183,258 base pairs and 19 to 172 genes.

Determining how removing large regions of a genome impacts viability and growth is a first step toward achieving a streamlined genome. The success of 19 of our 55 attempts to erase large regions from the *A. baylyi* genome under rich medium conditions depended to some extent on whether they contained genes judged to be essential for viability or maintaining high fitness when they were inactivated one-at-a-time. There was a higher correlation between deletion success for genes absolutely required for viability in a single-gene knockout collection in minimal medium than there was for genes that were needed to maintain high fitness in a Tn-Seq library constructed in rich medium. These single-gene results were not able to completely predict whether a larger deletion containing multiple genes would be viable, and the exceptions can give interesting information about how cellular components in ADP1 are organized and interact. For example, there can be synthetic lethals in which deletion of two or more genes leads to a loss in viability even when none of the genes is essential on their own (62, 63). Multiple-gene deletions may also synergistically perturb gene expression and metabolic pathways in ways that create deleterious imbalances that are bigger than for single-gene deletions (64).

Only 2 of the 19 *A. baylyi* large deletions lost their ability to grow in minimal medium. This result was expected for one of these strains (MGD15) because it should be incapable of pyrimidine biosynthesis due to deletion of the *pyrF* gene. Supplementation with uracil was able to restore growth of this deletion strain in minimal succinate medium, showing that *pyrF* could be used as an auxotrophic selection marker in ADP1 as it is in many other bacterial species (65). The other such deletion (MGD9) did not contain any genes that led to inviability in minimal medium when they were deleted one at a time in the knockout strain collection. ADP1 encodes two isocitrate dehydrogenase isozymes encoded by the *ACIAD1187* and *ACIAD1190* genes (30). Both of these genes were simultaneously deleted in this strain. We suspect that this leads to synthetic lethality in this minimal medium that has succinate as the sole carbon source.

Interestingly, 4 of the 19 MGD strains exhibited growth in minimal medium despite the removal of genes that were found to be essential in the single-gene knockout collection. MGD8 included the deletion of four putative essential genes (*lysS*, *cysD*, *cysN* and ACIAD1056). MGD13 showed growth despite lacking two such genes (*terD* and *ACIAD1965*). MGD7 deleted an ATP-binding subunit of an iron transporter (*ACIAD0969*) and a gene of unknown function (*ACIAD1000*) that were essential in minimal medium. Finally, MGD19 included a putative gene of unknown function (*ACIAD3600*) essential for growth in minimal medium.

Many successful deletions—in 9 of the 19 MGD strains—included genes found to be essential for fitness in the Tn-Seq experiment in rich medium that were not essential for viability in minimal medium. These genes included several sugar transferases (*ACIAD0084*-*ACIAD0091*), a lipid metabolism gene (*fadB*), an outer membrane protein (*ACIAD0697*), an elongation factor (*typA*), a putative hemolysin (*ACIAD0944*), isocitrate dehydrogenase (*idh*), a putative transthyretin-like protein (*ACIAD1188*), a succinylglutamate desuccinylase (*astE*), a transcriptional regulator (*dcaS*), an efflux pump(*smvA*), a uridylyltransferase (*glnD*), and cysteine synthase A (*cysK*), another putative transcriptional regulator (ACIAD1082) and a penicillin-binding protein (*pbpA*). Loss of these genes may contribute to the reduced growth rates of the multiple-gene deletion strains, making them top candidates for adding back one or a few of the genes removed in each segment to maintain high fitness.

The overall worse growth of the MGD strains may be the result from metabolic imbalances that result from genome reduction (66) or deletion of quasi-essential genes (18). The deletions most detrimental to growth were in the MGD15 and MGD9 strains. A possible explanation is that among the genes deleted in MGD15 is a gene essential in succinate minimal media (*pyrF*) in addition to malate synthase G (*glcB*), a gene involved in the TCA cycle which is essential for growth in acetate and 2,3-butanediol as sole carbon sources (53). Similarly, two genes deleted in MGD9, *ACIAD2740* and *ACIAD2741*, were previously determined to be critical for growth in acetate and quinate, respectively (53) and their loss may be particularly disruptive to metabolism.

Some of the engineered MGDs may retain reasonable growth rates because the effects of deleting different genes balance one another out. For instance, MGD1 may emulate mutations inactivating the *per* and *pgi* genes that have been observed in an adaptive laboratory evolution experiment with ADP1 (46). The beneficial deletion of these genes may balance the detrimental effects generated by the rest of the content in the deletion. In this case, the inclusion of *galU* in MGD1 could be counteracting the fitness-enhancing effects of *per* and *pgi* deletions, since a *galU* knockout shows defective growth in acetate, 2,3-butanediol, and quinate (53). MGD1 may also have benefited from the removal of an entire fimbrial operon (*ACIAD0119*-*ACIAD0123*), as these structures are often costly to express and have no benefit under lab conditions. Similarly, MGD5 may have enhanced fitness from deleting the type IV fimbrial biogenesis gene *fimT*.

Other deletions may illustrate the advantages and disadvantages of concentrating on genome minimization in one environment while not examining robustness to other environments. MGD5 includes several possible detrimental deletions: a stress gene *ACIAD0704* (*mscL* homolog with known functions in hypoosmotic shock and in mechanical, nutritional and oxidative stress), DNA repair gene *mutM* (formamidopyrimidine-DNA glycosylase) and *tonB* (heme/iron acquisition). It is likely that this deletion strain is less robust in terms of both its genome stability and its ability to survive nutrient and physical stresses. MGD11, one of the larger deletions (183 kb and 167 genes), tolerates the somewhat surprising elimination of one copy of the 16S and 23S ribosomal RNA genes, along with Ala and Ile tRNA copies. While it does not delete any genes that are essential for viability in minimal media, it does remove two genes that are essential for fitness in LB, *cysK* and *dcaS*. In this case, it is possible that there could be a fitness balancing effect of simultaneously deleting multiple pathways involved in the degradation of aromatic compounds including the *pca*-*qui*-*pob*-*hca* gene cluster (28, 32).

Overall, we found that the effect of simultaneously deleting many genes on viability/fitness cannot be fully predicted from the impact of single-gene deletions. Computational methods that use genome-scale models, which can account for certain types of interactions between genes and pathways, have the potential to better inform the deletion process (19), including how one can relocate of one or a few genes that are needed for viability or fitness from a stretch of chromosome before it is deleted. The *pyrH* and *icd* genes are candidates for this approach. They could be added back in place of the deleted region to the respective MGD strains during the Golden Transformation rescue step. Perhaps a bigger issue for the genome reduction enterprise is the potential for losing robustness in other environments when genes are deleted from a strain. The extent of this loss of robustness remains to be tested for the MGD strains in future work.

RNA-guided nucleases have revolutionized the field of genome editing, among many other applications. Much of this work has focused on CRISPR-Cas9, but there are a large diversity of CRISPR types in bacterial genomes that are less well-characterized (67). The type I-Fa CRISPR-Cas system of *A. baylyi* is analogous to the type I-Fb CRISPR/Cas system of *A. baumanii*, except the *cas1* gene appears last in the Cas operon (58, 60). *A. baylyi* ADP1 contains three CRISPR arrays: one with 91 spacers adjacent to the Cas operon and two others with 21 and 6 spacers, respectively, located elsewhere in the genome. By using Golden Transformation to replace the 91-spacer CRISPR array with the *tdk*-*kanR* dual selection cassette to create a “CRISPR-Ready” strain and then inserting a single synthetic spacer in place of this cassette, we showed that *A. baylyi*’s CRISPR system is active and readily reprogrammable.

Novel spacers may be designed and transformed into a CRISPR-Ready *A. baylyi* strain, allowing this system to be repurposed for various applications. As an example, we created a “self-targeting” CRISPR-Lock that tests whether a certain gene has been successfully removed from the chromosome and prevents its re-acquisition. The large deletions in the strains we created could be combined in future work by adding genomic DNA isolated from one deletion strain to a locked version of another deletion strain. This strategy could be particularly effective if self-targeting spacers are added in place of each deleted region during the rescue step that removed the *tdk*-*kanR* cassette so that this protection will also accumulate during genome streamlining. Although orthogonal dCas9 gene repression has been applied successfully in *A. baylyi* ADP1, there was a metabolic cost for expressing this foreign system (68). The ability to reprogram the endogenous CRISPR/Cas type I-F system opens up the possibility of employing it for transcriptional repression (analogous to dCas9) via *cas3* knockout (69) without this burden issue. Repurposing the native CRISPR system in this way would provide a new tool for implementing genome-wide screens and combinatorial pathway engineering in this strain.

In summary, we demonstrated new methods for more facile and assured genome editing in *Acinetobacter baylyi*. We used Golden Transformation to create 19 large deletions that outline a future roadmap for significantly reducing its genome complexity to make it a simpler and more predictable metabolic and genome engineering platform. This method also simplifies assembling multiple DNA parts in a one-pot reaction before they are added to the genome, opening up new possibilities for constructing and testing combinatorial libraries. Additionally, by showing that the native *A. baylyi* CRISPR-Cas system is active we also create further avenues for restricting transformation of DNA that might reverse our engineered deletions and for repurposing this system for genetic control. These developments leverage and further improve upon *A. baylyi*’s remarkable utility as a chassis organism that is highly naturally transformable.

## DATA AVAILABILITY

Genome sequencing reads have been deposited with the National Center for Biotechnology Information Sequence Read Archive under project accession number PRJNA559727.

## ACKNOWLEDGEMENT

We thank Ellen Neidle for sharing the puddle transformation approach.

## REFERENCES

1. Arkin, A.P. and Fletcher, D.A. (2006) Fast, cheap and somewhat in control. Genome Biol., 7, 114.

2. Renda, B.A., Hammerling, M.J. and Barrick, J.E. (2014) Engineering reduced evolutionary potential for synthetic biology. Mol. Biosyst., 10, 1668–1678.

3. Juhas, M., Reuß, D.R., Zhu, B. and Commichau, F.M. (2014) *Bacillus subtilis* and *Escherichia coli* essential genes and minimal cell factories after one decade of genome engineering. Microbiol. (United Kingdom), 160, 2341–2351.

4. Calero, P. and Nikel, P.I. (2019) Chasing bacterial chassis for metabolic engineering: a perspective review from classical to non-traditional microorganisms. Microb. Biotechnol., 12, 98–124.

5. Fehér, T., Papp, B., Pal, C. and Pósfai, G. (2007) Systematic genome reductions: theoretical and experimental approaches. Chem. Rev., 107, 3498–3513.

6. Leprince, A., van Passel, M.W.J. and dos Santos, V.A.P.M. (2012) Streamlining genomes: toward the generation of simplified and stabilized microbial systems. Curr. Opin. Biotechnol., 23, 651–658.

7. Glass, J.I., Merryman, C., Wise, K.S., Hutchison, C.A. and Smith, H.O. (2017) Minimal cells— Real and imagined. Cold Spring Harb. Perspect. Biol., 9, a023861.

8. Martínez-García, E. and de Lorenzo, V. (2016) The quest for the minimal bacterial genome. Curr. Opin. Biotechnol., 42, 216–224.

9. Park, M.K., Lee, S.H., Yang, K.S., Jung, S.C., Lee, J.H. and Kim, S.C. (2014) Enhancing recombinant protein production with an *Escherichia coli* host strain lacking insertion sequences. Appl. Microbiol. Biotechnol., 98, 6701–6713.

10. Pósfai, G., Plunkett, G., Fehér, T., Frisch, D., Keil, G.M., Umenhoffer, K., Kolisnychenko, V., Stahl, B., Sharma, S.S., de Arruda, M., et al. (2006) Emergent properties of reduced-genome *Escherichia coli*. Science, 312, 1044–1046.

11. Suárez, G.A., Renda, B.A., Dasgupta, A. and Barrick, J.E. (2017) Reduced mutation rate and increased transformability of transposon-free *Acinetobacter baylyi* ADP1-ISx. Appl. Environ. Microbiol., 83, e01025–17.

12. Choi, J.W., Yim, S.S., Kim, M.J. and Jeong, K.J. (2015) Enhanced production of recombinant proteins with *Corynebacterium glutamicum* by deletion of insertion sequences (IS elements). Microb. Cell Fact., 14, 207.

13. Martínez-García, E., Nikel, P.I., Aparicio, T. and de Lorenzo, V. (2014) *Pseudomonas* 2.0: genetic upgrading of *P. putida* KT2440 as an enhanced host for heterologous gene expression. Microb. Cell Fact., 13, 159.

14. Martínez-García, E., Nikel, P.I., Chavarría, M. and de Lorenzo, V. (2014) The metabolic cost of flagellar motion in *Pseudomonas putida* KT2440. Environ. Microbiol., 16, 291–303.

15. Komatsu, M., Uchiyama, T., Omura, S., Cane, D.E. and Ikeda, H. (2010) Genome-minimized *Streptomyces* host for the heterologous expression of secondary metabolism. Proc. Natl. Acad. Sci. U. S. A., 107, 2646–2651.

16. Wang, Z., Huang, X., Hu, H., Wang, W., Shen, X. and Zhang, X. (2017) Developing genome-reduced *Pseudomonas chlororaphis* strains for the production of secondary metabolites. BMC Genomics, 18, 715.

17. Mizoguchi, H., Sawano, Y., Kato, J.I. and Mori, H. (2008) Superpositioning of deletions promotes growth of *Escherichia coli* with a reduced genome. DNA Res., 15, 277–284.

18. Hutchison, C.A., Chuang, R.-Y., Noskov, V.N., Assad-Garcia, N., Deerinck, T.J., Ellisman, M.H., Gill, J., Kannan, K., Karas, B.J., Ma, L., et al. (2016) Design and synthesis of a minimal bacterial genome. Science, 351, aad6253.

19. Wang, L. and Maranas, C.D. (2018) MinGenome: An *in silico* top-down approach for the synthesis of minimized genomes. ACS Synth. Biol., 7, 462–473.

20. Kurokawa, M., Seno, S., Matsuda, H. and Ying, B.-W. (2016) Correlation between genome reduction and bacterial growth. DNA Res., 23, 517–525.

21. Hashimoto, M., Ichimura, T., Mizoguchi, H., Tanaka, K., Fujimitsu, K., Keyamura, K., Ote, T., Yamakawa, T., Yamazaki, Y., Mori, H., et al. (2005) Cell size and nucleoid organization of engineered *Escherichia coli* cells with a reduced genome. Mol. Microbiol., 55, 137–149.

22. Rancati, G., Moffat, J., Typas, A. and Pavelka, N. (2018) Emerging and evolving concepts in gene essentiality. Nat. Rev. Genet., 19, 34–49.

23. Metzgar, D., Bacher, J.M., Pezo, V., Reader, J., Döring, V., Schimmel, P., Marlière, P. and de Crécy-Lagard, V. (2004) Acinetobacter sp. ADP1: an ideal model organism for genetic analysis and genome engineering. Nucleic Acids Res., 32, 5780–5790.

24. Elliott, K.T. and Neidle, E.L. (2011) *Acinetobacter baylyi* ADP1: transforming the choice of model organism. IUBMB Life, 63, 1075–1080.

25. Seaton, S.C., Elliott, K.T., Cuff, L.E., Laniohan, N.S., Patel, P.R. and Neidle, E.L. (2012) Genome-wide selection for increased copy number in *Acinetobacter baylyi* ADP1: locus and context-dependent variation in gene amplification. Mol. Microbiol., 83, 520–535.

26. Renda, B.A., Chan, C., Parent, K.N. and Barrick, J.E. (2016) Emergence of a competence-reducing filamentous phage from the genome of *Acinetobacter baylyi* ADP1. J. Bacteriol., 198, 3209–3219.

27. Leong, C.G., Bloomfield, R.A., Boyd, C.A., Dornbusch, A.J., Lieber, L., Liu, F., Owen, A., Slay, E., Lang, K.M. and Lostroh, C.P. (2017) The role of core and accessory type IV pilus genes in natural transformation and twitching motility in the bacterium *Acinetobacter baylyi*. PLoS ONE, 12, e0182139.

28. Young, D.M., Parke, D. and Ornston, L.N. (2005) Opportunities for genetic investigation afforded by *Acinetobacter baylyi*, a nutritionally versatile bacterial species that is highly competent for natural transformation. Annu. Rev. Microbiol., 59, 519–551.

29. Fischer, R., Bleichrodt, F.S. and Gerischer, U.C. (2008) Aromatic degradative pathways in *Acinetobacter baylyi* underlie carbon catabolite repression. Microbiology, 154, 3095–3103.

30. Barbe, V., Vallenet, D., Fonknechten, N., Kreimeyer, A., Oztas, S., Labarre, L., Cruveiller, S., Robert, C., Duprat, S., Wincker, P., et al. (2004) Unique features revealed by the genome sequence of Acinetobacter sp. ADP1, a versatile and naturally transformation competent bacterium. Nucleic Acids Res., 32, 5766–5779.

31. de Berardinis, V., Durot, M., Weissenbach, J. and Salanoubat, M. (2009) *Acinetobacter baylyi* ADP1 as a model for metabolic system biology. Curr. Opin. Microbiol., 12, 568–576.

32. Stuani, L., Lechaplais, C., Salminen, A. V., Ségurens, B., Durot, M., Castelli, V., Pinet, A., Labadie, K., Cruveiller, S., Weissenbach, J., et al. (2014) Novel metabolic features in *Acinetobacter baylyi* ADP1 revealed by a multiomics approach. Metabolomics, 10, 1223–1238.

33. Lehtinen, T., Efimova, E., Santala, S. and Santala, V. (2018) Improved fatty aldehyde and wax ester production by overexpression of fatty acyl-CoA reductases. Microb. Cell Fact., 17, 19.

34. Santala, S., Efimova, E., Kivinen, V., Larjo, A., Aho, T., Karp, M. and Santala, V. (2011) Improved triacylglycerol production in *Acinetobacter baylyi* ADP1 by metabolic engineering. Microb. Cell Fact., 10, 36.

35. Kok, R.G., D’Argenio, D.A. and Ornston, L.N. (1997) Combining localized PCR mutagenesis and natural transformation in direct genetic analysis of a transcriptional regulator gene, *pobR*. J. Bacteriol., 179, 4270–4276.

36. Jones, R.M. and Williams, P.A. (2003) Mutational analysis of the critical bases involved in activation of the AreR-regulated σ^54^-dependent promoter in *Acinetobacter* sp. strain ADP1. Appl. Environ. Microbiol., 69, 5627–5635.

37. Tumen-Velasquez, M., Johnson, C.W., Ahmed, A., Dominick, G., Fulk, E.M., Khanna, P., Lee, S.A., Schmidt, A.L., Linger, J.G., Eiteman, M.A., et al. (2018) Accelerating pathway evolution by increasing the gene dosage of chromosomal segments. Proc. Natl. Acad. Sci., 115, 7105–7110.

38. Engler, C., Kandzia, R. and Marillonnet, S. (2008) A one pot, one step, precision cloning method with high throughput capability. PLoS ONE, 3.

39. Lee, M.E., DeLoache, W.C., Cervantes, B. and Dueber, J.E. (2015) A highly characterized yeast toolkit for modular, multipart assembly. ACS Synth. Biol., 4, 975–986.

40. R Core Team (2019) R: A Language and Environment for Statistical Computing R Foundation for Statistical Computing, Vienna, Austria.

41. Kulasekara, H.D., Ventre, I., Kulasekara, B.R., Lazdunski, A., Filloux, A. and Lory, S. (2005) A novel two-component system controls the expression of *Pseudomonas aeruginosa* fimbrial *cup* genes. Mol. Microbiol., 55, 368–380.

42. Demarre, G., Guérout, A.M., Matsumoto-Mashimo, C., Rowe-Magnus, D. a., Marlière, P. and Mazel, D. (2005) A new family of mobilizable suicide plasmids based on broad host range R388 plasmid (IncW) and RP4 plasmid (IncPα) conjugative machineries and their cognate *Escherichia coli* host strains. Res. Microbiol., 156, 245–255.

43. Powell, J.E., Leonard, S.P., Kwong, W.K., Engel, P. and Moran, N.A. (2016) Genome-wide screen identifies host colonization determinants in a bacterial gut symbiont. Proc. Natl. Acad. Sci. U. S. A., 113, 13887–13892.

44. Love, M.I., Huber, W. and Anders, S. (2014) Moderated estimation of fold change and dispersion for RNA-seq data with DESeq2. Genome Biol., 15, 550.

45. Huerta-Cepas, J., Forslund, K., Coelho, L.P., Szklarczyk, D., Jensen, L.J., Von Mering, C. and Bork, P. (2017) Fast genome-wide functional annotation through orthology assignment by eggNOG-mapper. Mol. Biol. Evol., 34, 2115–2122.

46. Renda, B.A., Dasgupta, A., Leon, D. and Barrick, J.E. (2015) Genome instability mediates the loss of key traits by Acinetobacter baylyi ADP1 during laboratory evolution. J. Bacteriol., 197, 872–881.

47. Deatherage, D.E. and Barrick, J.E. (2014) Identification of mutations in laboratory-evolved microbes from next-generation sequencing data using *breseq*. Methods Mol. Biol., 1151, 165–188.

48. Barrick, J.E., Colburn, G., Deatherage, D.E., Traverse, C.C., Strand, M.D., Borges, J.J., Knoester, D.B., Reba, A. and Meyer, A.G. (2014) Identifying structural variation in haploid microbial genomes from short-read resequencing data using *breseq*. BMC Genomics, 15, 1039.

49. Murin, C.D., Segal, K., Bryksin, A. and Matsumura, I. (2012) Expression vectors for *Acinetobacter baylyi* ADP1. Appl. Environ. Microbiol., 78, 280–283.

50. Overballe-Petersen, S., Harms, K., Orlando, L. a. a., Mayar, J.V.M., Rasmussen, S., Dahl, T.W., Rosing, M.T., Poole, A.M., Sicheritz-Ponten, T., Brunak, S., et al. (2013) Bacterial natural transformation by highly fragmented and damaged DNA. Proc. Natl. Acad. Sci. U. S. A., 110, 19860–19865.

51. Simpson, D., Dawson, L. and Fry, J. (2007) Influence of flanking homology and insert size on the transformation frequency of *Acinetobacter baylyi* BD413. Environ. Biosafety Res., 6, 55–69.

52. Leonard, S.P., Perutka, J., Powell, J.E., Geng, P., Richhart, D.D., Byrom, M., Kar, S., Davies, B.W., Ellington, A.D., Moran, N.A., et al. (2018) Genetic engineering of bee gut microbiome bacteria with a toolkit for modular assembly of broad-host-range plasmids. ACS Synth. Biol., 7, 1279–1290.

53. de Berardinis, V., Vallenet, D., Castelli, V., Besnard, M., Pinet, A., Cruaud, C., Samair, S., Lechaplais, C., Gyapay, G., Richez, C., et al. (2008) A complete collection of single-gene deletion mutants of *Acinetobacter baylyi* ADP1. Mol. Syst. Biol., 4, 174.

54. Baba, T., Ara, T., Hasegawa, M., Takai, Y., Okumura, Y., Baba, M., Datsenko, K.A., Tomita, M., Wanner, B.L. and Mori, H. (2006) Construction of *Escherichia coli* K-12 in-frame, single-gene knockout mutants: the Keio collection. Mol. Syst. Biol., 2, 2006.0008.

55. Yamamoto, N., Nakahigashi, K., Nakamichi, T., Yoshino, M., Takai, Y., Touda, Y., Furubayashi, A., Kinjyo, S., Dose, H., Hasegawa, M., et al. (2009) Update on the Keio collection of *Escherichia coli* single-gene deletion mutants. Mol. Syst. Biol., 5, 335.

56. Goodall, E.C.A., Robinson, A., Johnston, I.G., Jabbari, S., Turner, K.A., Cunningham, A.F., Lund, P.A., Cole, J.A. and Henderson, I.R. (2018) The essential genome of *Escherichia coli* K-12. mBio, 9, e02096–17.

57. Reams, A.B. and Neidle, E.L. (2004) Gene amplification involves site-specific short homology-independent illegitimate recombination in *Acinetobacter* sp. strain ADP1. J. Mol. Biol., 338, 643–656.

58. Touchon, M., Cury, J., Yoon, E.-J., Krizova, L., Cerqueira, G.C., Murphy, C., Feldgarden, M., Wortman, J., Clermont, D., Lambert, T., et al. (2014) The genomic diversification of the whole *Acinetobacter* genus: origins, mechanisms, and consequences. Genome Biol. Evol., 6, 2866–2882.

59. Mojica, F.J.M., Díez-Villaseñor, C., García-Martínez, J. and Almendros, C. (2009) Short motif sequences determine the targets of the prokaryotic CRISPR defence system. Microbiology, 155, 733–740.

60. Karah, N., Samuelsen, Ø., Zarrilli, R., Sahl, J.W., Wai, S.N. and Uhlin, B.E. (2015) CRISPR-cas subtype I-Fb in *Acinetobacter baumannii*: evolution and utilization for strain subtyping. PLoS ONE, 10, e0118205.

61. Biswas, A., Gagnon, J.N., Brouns, S.J.J., Fineran, P.C. and Brown, C.M. (2013) CRISPRTarget: Bioinformatic prediction and analysis of crRNA targets. RNA Biol., 10, 817–827.

62. Mori, H., Baba, T., Yokoyama, K., Takeuchi, R., Nomura, W., Makishi, K., Otsuka, Y., Dose, H. and Wanner, B.L. (2015) Identification of essential genes and synthetic lethal gene combinations in *Escherichia coli* K-12. Methods Mol. Biol., 1279, 45–65.

63. Typas, A., Nichols, R.J., Siegele, D.A., Shales, M., Collins, S.R., Lim, B., Braberg, H., Yamamoto, N., Takeuchi, R., Wanner, B.L., et al. (2008) High-throughput, quantitative analyses of genetic interactions in *E. coli*. Nat. Methods, 5, 781–787.

64. Reuß, D.R., Altenbuchner, J., Mäder, U., Rath, H., Ischebeck, T., Sappa, P.K., Thürmer, A., Guérin, C., Nicolas, P., Steil, L., et al. (2017) Large-scale reduction of the *Bacillus subtilis* genome: consequences for the transcriptional network, resource allocation, and metabolism. Genome Res., 27, 289–299.

65. Freed, E., Fenster, J., Smolinski, S.L., Walker, J., Henard, C.A., Gill, R. and Eckert, C.A. (2018) Building a genome engineering toolbox in nonmodel prokaryotic microbes. Biotechnol. Bioeng., 115, 2120–2138.

66. Choe, D., Lee, J.H., Yoo, M., Hwang, S., Sung, B.H., Cho, S., Palsson, B., Kim, S.C. and Cho, B.-K. (2019) Adaptive laboratory evolution of a genome-reduced *Escherichia coli*. Nat. Commun., 10, 935.

67. Makarova, K.S., Haft, D.H., Barrangou, R., Brouns, S.J.J., Charpentier, E., Horvath, P., Moineau, S., Mojica, F.J.M., Wolf, Y.I., Yakunin, A.F., et al. (2011) Evolution and classification of the CRISPR-Cas systems. Nat. Rev. Microbiol., 9, 467–477.

68. Geng, P., Leonard, S.P., Mishler, D.M. and Barrick, J.E. (2019) Synthetic genome defenses against selfish DNA elements stabilize engineered bacteria against evolutionary failure. ACS Synth. Biol., 8, 521–531.

69. Luo, M.L., Mullis, A.S., Leenay, R.T. and Beisel, C.L. (2015) Repurposing endogenous type I CRISPR-Cas systems for programmable gene repression. Nucleic Acids Res., 43, 674–681.

