## Supplementary material for "Rapid and assured genetic engineering methods applied to *Acinetobacter baylyi* ADP1 genome streamlining": Figure S1

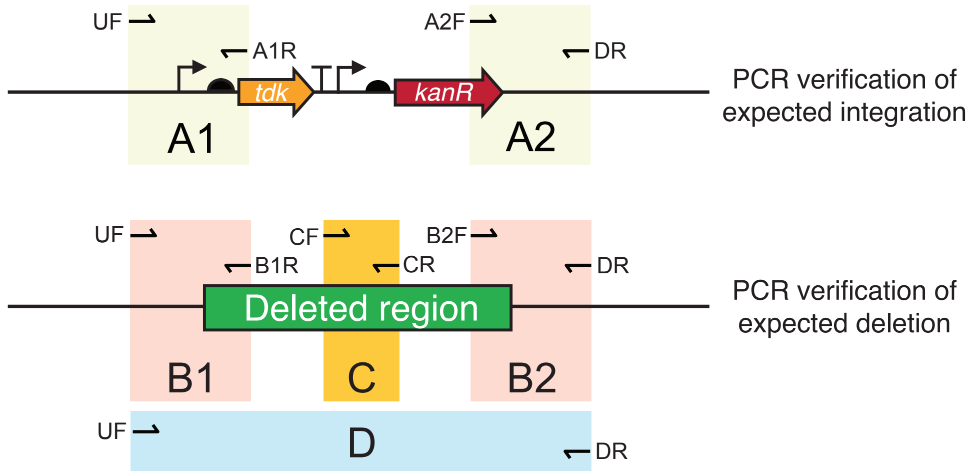


**Figure S1. PCR reactions used to validate multiple-gene deletion strain construction.** PCR reactions A1 and A2 check for insertion of the *tdk-kanR* cassette in the proper context. PCR reactions B1, B2, C, and D verify that the deleted region is missing from the edited chromosome. **Table S1** lists the primer sequences used in each reaction for each designed ADP1 deletion. **Table S3** reports the results of each type of PCR reaction for each candidate deletion strain.
